# Sex-specific glycosylation of secreted immunomodulatory proteins in the filarial nematode *Brugia malayi*

**DOI:** 10.1101/2021.02.24.432741

**Authors:** Joseph Koussa, Burcu Vitrinel, Peter Whitney, Brian Kasper, Lara K. Mahal, Christine Vogel, Sara lustigman, Kourosh Salehi-Ashtiani, Elodie Ghedin

## Abstract

The extended persistence of filarial nematodes within a host suggests immunomodulatory mechanisms that allow the parasites to resist or evade the host immune response. There is increasing evidence for immunomodulatory glycans expressed by a diversity of parasitic worms. In this study, we integrate multiple layers of the host-parasite interface to investigate the glycome of a model filarial parasite, *Brugia malayi*. We report a significant overrepresentation of terminal GalNAc moieties in adult female worms coupled with an overall upregulation in O-glycosylation, T-antigen expression, and a bias for galactose containing glycans. Adult males preferentially displayed a bias for terminal GlcNAc containing glycans, and fucosylated epitopes. Subsequent proteomic analysis confirmed sex-biases in protein glycosylation and highlighted the sex-specific glycosylation of well characterized immunomodulators expressed and secreted by *B. malayi*. We identify sex-specific effectors at that interface and suggest approaches to selectively interfere with the parasitic life cycle and potentially control transmission.

## Introduction

*Brugia malayi* is a parasitic nematode and, along with *Wuchereria bancrofti*, a causative agent of lymphatic filariasis in humans. The life cycle of *B. malayi* involves both a mosquito vector and a human host in which the worms can persist for up to eight years as adults colonizing the lymphatic system (Nutman). In contrast to the majority of nematodes, including the free-living model nematode *C. elegans*, parasitic worms like *B. malayi* are sexually dimorphic, with male and female worms coexisting within the same host. Once sexual maturity is reached (>120 days post infection), fertilized adult females release microfilariae (mf)—the youngest life stage of the worm—into blood circulation. These are then picked up by the mosquito during a blood meal. In the mosquito, the mf develops into infective stage 3 larvae (L3), which can be transmitted to a new human host during a subsequent blood meal.

Sexual dimorphism in parasitic nematodes has been suggested to be a response to host immune pressure (Gemmill et al.). Sexual dimorphism has been well characterized in *B. malayi* (Michalski and Weil; Jiang, Li, et al.; Kashyap et al.). At both the transcriptomic and proteomic levels, adult male and female *B. malayi* display sex biases in gene expression with adult males characterized by enrichment for genes involved in energy production, metabolic processes and cytoskeletal proteins, while adult females have gene expression profiles enriched in signatures for RNA modification and transcription (Jiang, Malone, et al.). Proteomic analysis of sex-specific secretomes from *B. malayi* identified significant differences between adult male and female worms; 70% of proteins secreted by adult males and 65% secreted by adult females were unique to each sex (Bennuru et al.).

More recently, proteomic analysis of exosome-like vesicles secreted from adult *B. malayi* revealed sex-dependence in protein cargo. Adult female-secreted vesicles had a high content in Bma-Galectin-2, Triose Phosphate isomerase (TPI), Macrophage migration inhibitory factor (MIF1) and Thioredoxin peroxidase 2 among others, while the male-secreted vesicles were enriched for small GTPases, structural actin and tubulin, and a subset of heat shock proteins (Harischandra et al.). While sex differences in *B. malayi* have been investigated at both transcriptomic and proteomic levels, they remain understudied at the glycomic level.

Well conserved across evolution, glycosylation is a dynamic, non-templated process, dependent on enzymatic and substrate availability and resulting in a diversity of carbohydrate structures. Glycosylation plays key roles in regulating diverse biological processes, ranging from signaling and immune activation to development and reproduction in eukaryotic organisms (Varki, Ajit; Cummings, R. D.; Esko, J. D.; Freeze, H. H.; Stanley, P.; Bertozzi, C. R.; Hart, G. W.; Etzler and E.). Glycosylation is also known to act as a checkpoint in the proper folding of proteins within secretory pathways, as a large portion of secreted proteins from eukaryotes are glycosylated. In helminthic infections, glycoconjugates are involved in the underlying immunomodulation of infected hosts (Prasanphanich et al.; Harn et al.; Van Vliet et al.; Khoo and Dell). Several such glycoconjugates are at the host-parasite interface. For example, parasite excreted/secreted (ES) glycoproteins were shown to actively modulate the host immune system by physically interacting with cell surface receptors on dendritic cells (DC) and macrophages (van den Berg et al.; Rodríguez et al.). Studies have shown that in *B. malayi*, intact glycan structures on secreted proteins are necessary for the induction of the characteristic Th2 immune responses (Tawill et al.). In the blood fluke *S. mansoni*, studies revealed significant sex differences in glycan structures where females mainly carried Galβ1-4GlcNAc (Type II LacNAc) and Galβ1-4(Fucα1-3)GlcNAc (LewisX) antennae structures, whereas in males GalNAcβ1-4GlcNAc (Lacdi-NAc; LDN) and GalNAcβ1-4(Fucα1-3)GlcNAc (LDN-F) were prevalent in N-glycans, suggesting differential effects on host responses (Wuhrer et al.). Studies of the parasitic worms *Fasciola hepatica*, *Oesophagostomum dentatum*, and *S. mansoni* report sex and life stage biases in N-glycosylation profiles (Rodríguez et al.; Jiménez-Castells et al.). However, little information exists with respect to O-glycosylation trends and an in-depth characterization of glycosylation in filarial nematodes is lacking. Unraveling the sex-specific glycocode in *B. malayi* not only furthers our understanding of the biology of sexes and parasitism in nematodes, but also uncovers a set of new targets for anti-helminthic therapies or vaccine development, with the unique feature of potentially targeting male or female worms independently.

Herein we profile protein glycosylation in *B. malayi* using lectin microarray technology (Pilobello, Slawek, et al.; Agrawal et al.; Heindel et al.) and combine that with an expanded analysis of the sex-specific secretomes. We identified sex-dependent glycan and glycoprotein expression. We focused on the identification of sex-specific differentially glycosylated proteins at the host-parasite interface and evaluated their potential role in the immunomodulatory arsenal of *B. malayi*. We report the differential glycosylation in male and female *B. malayi* of three previously characterized immunomodulatory proteins, Bma-MIF-1 (Prieto-Lafuente et al.), Bma-FAR-1(Zhan et al.) and Bma-IPGM-1 (Singh et al.). We also report the sex-dependent glycosylation of two recently suggested drug targets, a phosphoglycerate kinase (Bm13839) (Kumar et al.) and a Calumenin (Bm5089) (Choi et al.), along with a subset of immune-relevant proteins, further suggesting sex-specific host-parasite interactions, that may be moderated by glycans, in *B. malayi* infections.

## Results

### *B. malayi* adult females secrete a larger diversity of proteins than adult males

Although sex-dependent gene expression in *B. malayi* worms has been well characterized (Bennuru et al.; Hewitson et al.; Grote et al.; Moreno and Geary), the stage- and sex-specific secretomes remain only partially defined. To gain a more complete understanding of the *B. malayi* secretome, in light of updated genome annotations and updated chromosome assemblies (Fauver et al, Tracey et al, Ghedin et al), we collected ES proteins from worms cultured *in vitro* at different life stages—mf, L3, L4, adult males, and adult females **(Fig. 1A).** Label-free mass spectrometric analysis of these samples identified a total of 1,114 unique proteins across life stages and sexes in the *B. malayi* secretome (**Supp. Table S1a**). These included previously identified proteins as well as an additional 444 proteins. It is worth noting that 47% of all identified proteins have no associated known functions. A summary table detailing overlaps with all preceding *B. malayi* proteomic studies can be found in **Supp. Table S1b.** We recovered 60% of proteins identified in (Mersha et al.) as GPI-anchored and 80% of proteins from exosome-like vesicles (EV) secreted by both adult sexes (Harischandra et al.). We also recovered 72% and 78% of proteins identified by Moreno et al and Hewiston et al, respectively, as part of *B. malayi’s* secretome (Moreno and Geary; Hewitson et al.). However, we only overlapped with 18% of the proteins identified by Bennuru et al (Bennuru et al.) as part of the secretome. Our data indicate that 70% of detected proteins were shared by at least two of the stages and that adult females secreted the largest diversity of proteins with 238 unique to this stage compared to 6 unique to adult males (**Fig. 1B)**.

**Figure 1:**
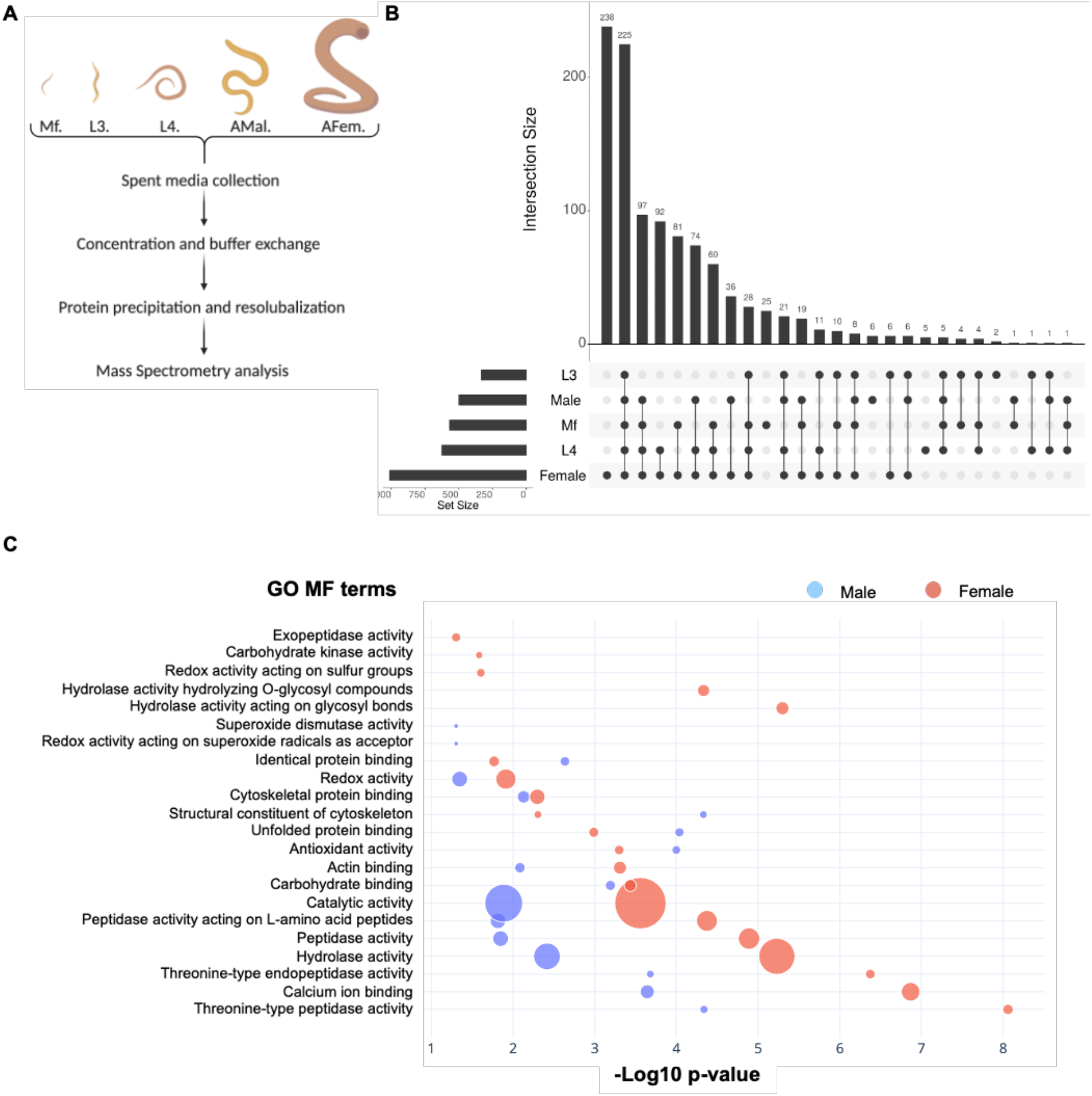
Proteomic characterization of stage- and sex-specific secretomes in *B. malayi*. **A.** Schematic representation of how *B. malayi* secretomes were collected for MS-MS analysis. **B.** UpsetR plot representing the intersections in stage- and sex-specific secretomes. The total number of proteins identified from each stage or sex are shown as set sizes. **C**. GO Molecular Function (MF) enrichment analysis of secreted proteins from adult males and females. Values shown represent–Log10 p-value and dot sizes are proportional to the number of proteins associated with each term.

To determine the functional implications of differentially secreted proteins, we performed functional enrichment on the sex-dependent protein sets (**Fig. 1C)**. In adult female worms, we observed specific enrichment of hydrolases acting on glycosyl residues, carbohydrate kinases and proteins involved in sulfur redox reactions (p-values < 0.05), while in adult males we see a significant enrichment of proteins involved in superoxide redox reactions and superoxide dismutase activities (p-values < 0.05). Male and female adult *B. malayi* secrete a large number of proteins with catalytic activity (e.g. threonine-type endopeptidases) and carbohydrate binding proteins, both of which are associated with helminth immunomodulatory properties (McSorley et al.).

### *B. malayi* has a glycogenome conserved across the filariae

As observed for most eukaryotic organisms, helminths glycosylate secreted proteins. These glycoproteins play key roles in immune function, and glycosylation is crucial to their activity (Cvetkovic et al.; Ahmed et al.). It is therefore important to define glycan structures decorating secreted proteins at the host-parasite interface. Determining the diversity of glycan structures expressed by an organism is largely dictated by the nature and number of glycosylation-related enzymes and available substrates. To construct a comprehensive list of glycosylation enzymes, we interrogated the *B. malayi* genome using a series of functional annotations. Gene Ontology (GO) and KEGG annotations for genes and pathways were used to assign glycosylation-related proteins. We identified 116 genes with an associated GO term relevant to protein glycosylation and 143 genes assigned to KEGG glycosylation pathways. A full BLASTp search of the *B. malayi* proteome against a “human glycosylation proteome” uncovered 136 genes with significant similarities. A full genome Hidden Markov Model scan for protein families (Pfam) was done to complement the analysis, uncovering 112 genes with Pfam annotations relevant to protein glycosylation. Overall, 285 genes were identified and were assigned to the *B. malayi* glycogenome (**Fig. 2A**; **Supp. Table S2a**).

**Figure 2:**
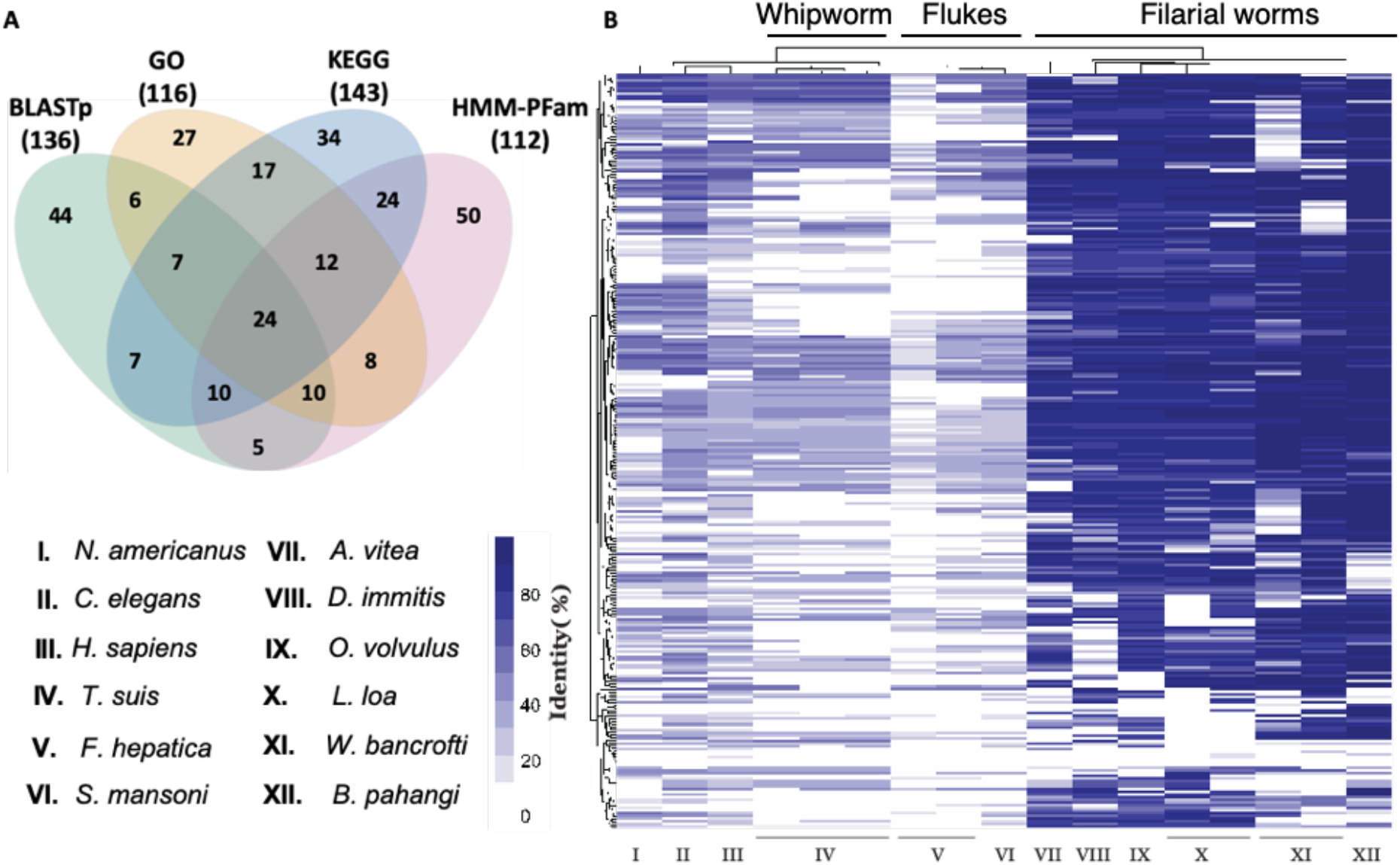
*B. malayi*’s glycogenome and ortholog analysis of glycosylation related genes across other worm genomes, and the human genome. **A.** Venn diagram representing the overlap analysis of the different functional annotation approaches used to identify glycosylation related genes in *B. malayi*. **B.** Ortholog analysis of all *B. malayi* glycosylation genes in parasitic and non-parasitic worms. Data shown represent the percentage in genetic identity between *B. malayi* glycosylation related genes and their corresponding orthologues in other organisms.

We evaluated the conservation of glycosylation across different clades of worms by comparative analysis of glycosylation-related orthologs in other helminth genomes. The analysis reveals a high degree of conservation within filarial nematodes (**Fig. 2B**, VII-XII) with around 56% of the genes conserved at >50% similarity. The data also indicated clear divergence from blood and liver flukes (**Fig. 2B**, V-VI) with 33% of the genes having no orthologs, and an intermediate profile of conservation when compared to hookworms, whipworms or the free-living nematode *C. elegans* with only 14% of the genes having no orthologs (**Fig. 2B**, I, IV, and II, respectively). Overall, the data show a higher conservation and similarity in the glycogenomes of filarial parasites and class III nematodes, and a clear divergence from trematodes such as *F. hepatica* and *S. mansoni*.

### *O*- and *N*-glycan biosynthetic pathways are differentially expressed between male and female adult worms

To investigate whether expression of the glycogenome is sex-dependent, we analyzed the transcriptome in adult male and female *B. malayi*. The overall expression profiles show high similarities between the sexes, as previously observed (Grote et al.). However, approximately 43% of the 285 glycosylation-related genes were significantly differentially expressed (p-value < 0.05). The majority of these genes belong to *O*-and *N*-glycan-related biosynthetic pathways. Female *B. malayi* significantly upregulate 50 glycosylation-related genes while males upregulate 72 (**Supp. Table S2b**). Mapping these results onto the biosynthetic pathways suggests higher expression of core 2/6 O-glycans in male worms (**Fig. 3A**). The results also indicate sex-dependent expression of specific fucose type O-glycans, with males having higher levels of GlcNAc-Fuc-(ser/thr), while females modify this epitope further to form predominantly Gal-GlcNAc-Fuc-(ser/thr) (**Fig. 3B**).

**Figure 3:**
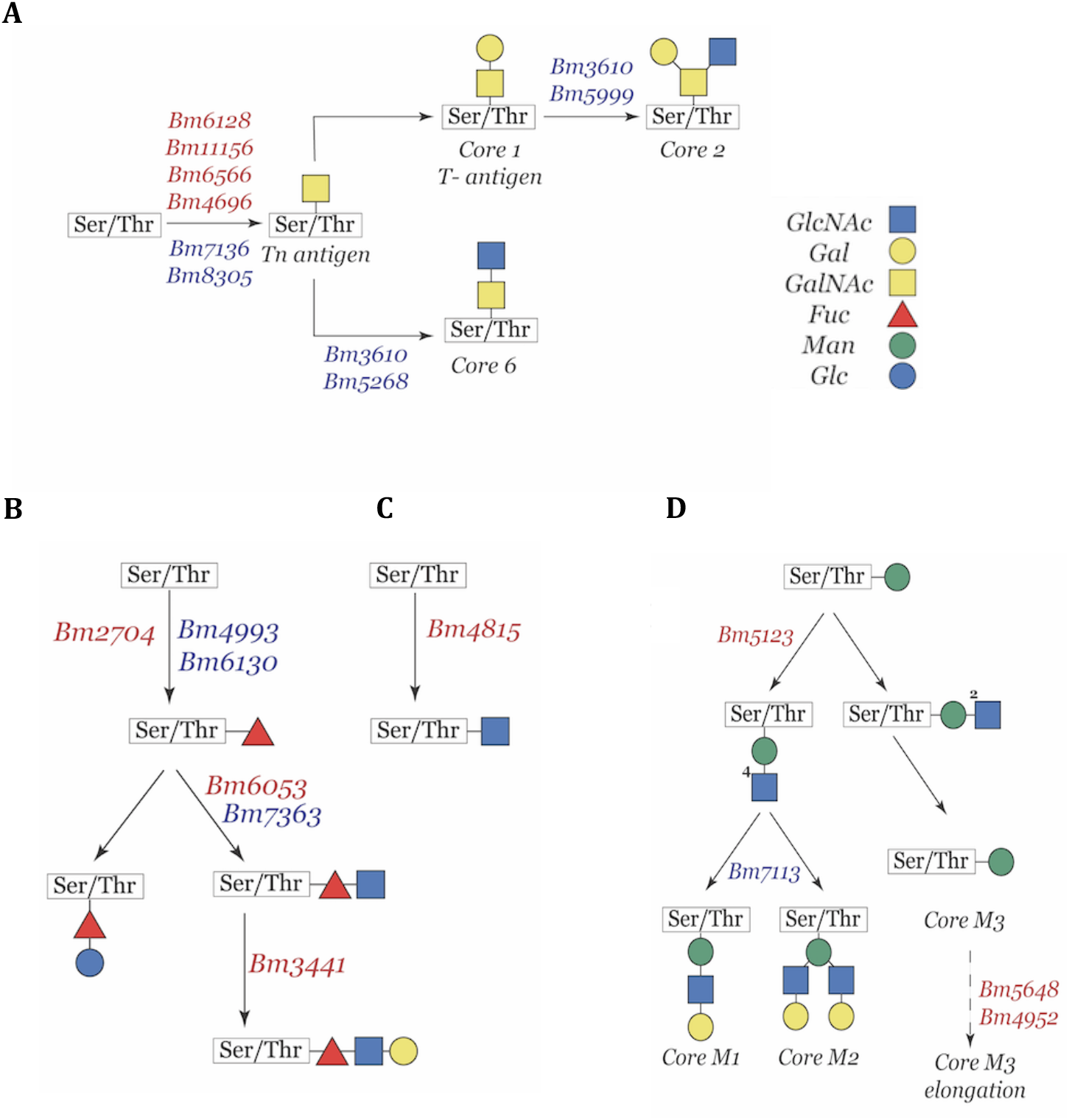
Differential expression of O-glycosylation related genes in adult male and female *B. malayi*. **A.** Partial representation of the mucin-type O-glycans biosynthetic pathway. *B. malayi* genes coding for enzymes catalyzing reactions are shown and color-coded. **B.** Fucose type O glycans biosynthetic pathway and corresponding *B. malayi* genes color-coded by upregulation status. **C.** N-acetylglucosamine type O-glycans biosynthetic pathway and corresponding *B. malayi* genes color-coded by upregulation status. **D.** Partial representation of the mannose type O-glycans biosynthetic pathway and corresponding *B. malayi* genes color-coded by upregulation status. Upregulated genes in females are highlighted in red and those upregulated in males are highlighted in blue.

Females also show a higher expression of a protein-O-acetylglucosamine transferase (Bm4815), leading to the uncommon GlcNAc O-linked glycan epitope (**Fig. 3C**). Similarly, the mannose type O-glycan biosynthetic pathway shows male-biased expression of core M1 and core M2 glycans (**Fig. 3D**). Sex-dependent expression profiles are also apparent for N-type glycan biosynthetic pathways where males display a higher expression of 17 genes within the N-type glycosylation pathway as compared to 7 genes with higher expression in females; most are N-glycan precursor and trimming enzymes. Both sexes also express genes related to glycan degradation, such as mannosidases and glycosidases, primarily involved in oligomannose and paucimannose N-glycan biosynthetic pathways (**Supp. Table S2c**).

### Glycan epitope expression is sex-biased in *B. malayi*

The study of glycosylation in *B. malayi* has primarily focused on GPI-anchored glycoproteins and staining of worms with specific lectins (Mersha et al.; Schraermeyer et al.; Kaushal et al.). A more general glycomic analysis of *B. malayi* has not been performed. To confirm whether the observed sex-biased differential expression of the *B. malayi* glycogenome translates into sex-biased display of glycan epitopes, we analyzed whole worm lysates on lectin microarrays (Pilobello, Krishnamoorthy, et al.; Propheter et al.). This technology uses the known glycan binding specificities of lectins to provide an epitope-specific readout of glycosylation. We observed significant differences in glycosylation profiles between male and female worms (**Fig. 4**). Female worms showed higher levels of terminal Gal/GalNAc epitopes, including core 1 O-glycans and T-antigen (lectins: MPA, MNA-G, HPA, ECA, GS-I, HAA, WFA, MNA-G, APP), and higher levels of LacDiNAc (LDN, lectin: SBA) and α1,2 fucosylation (TJA-II, PTL-II). In contrast, male worms had higher levels of terminal GlcNAc (lectins: WGA,) and fucosylated type 2 LacNAc epitopes (AAL).

**Figure 4:**
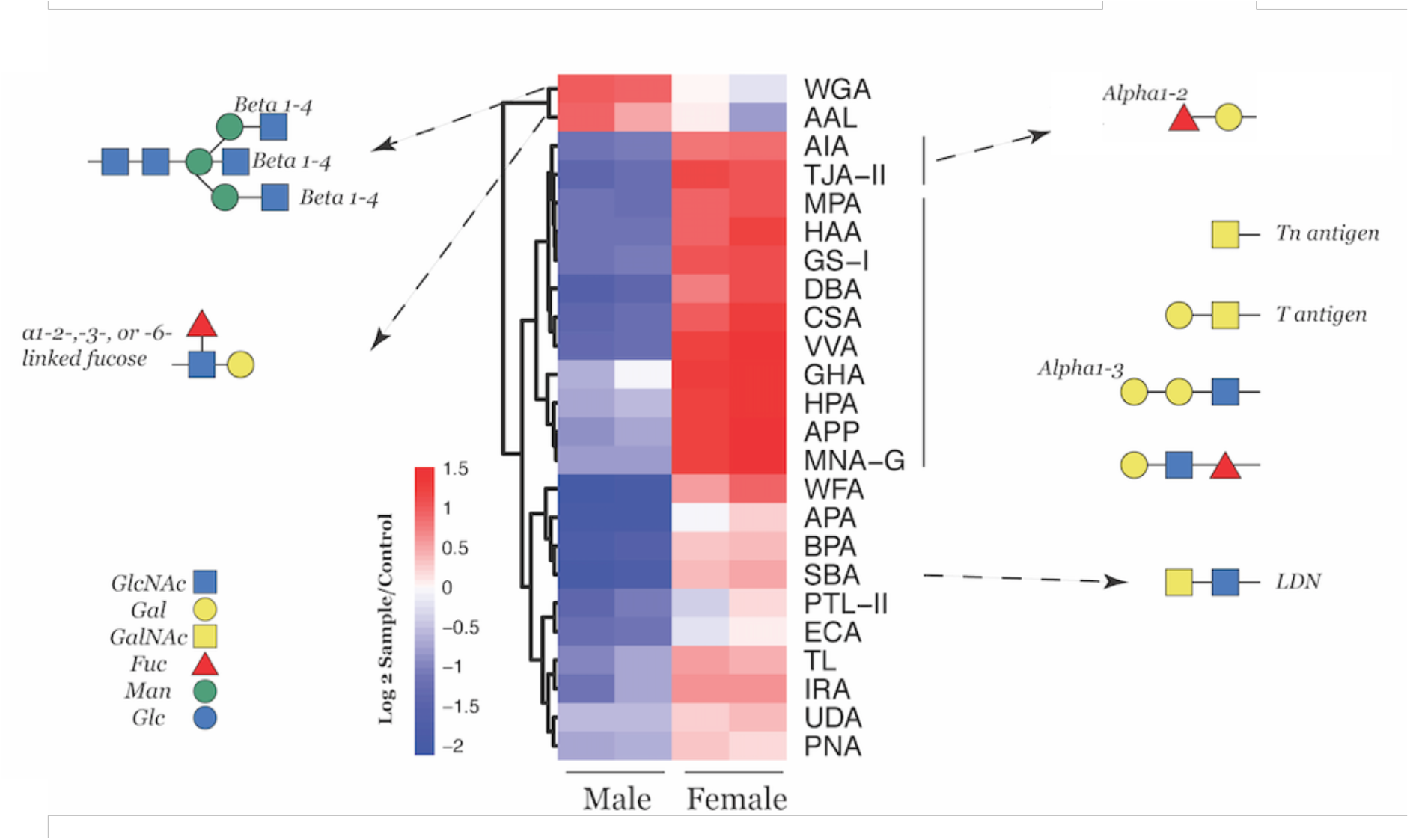
Glycomic profiling of *B. malayi* as determined by glycan binding to lectin arrays. Heatmap summarizing lectin array data comparing male and female glycans and representing significantly differentially displayed glycan epitopes between sexes (p < 0.05). Data shown represent Log2 of the fluorescence ratios between each sample and the control. Lectins are grouped by their corresponding recognition epitopes and representative glycan structure displaying respective epitopes are shown.

We suspected that biases in glycan epitopes would be associated with differential distribution within tissues in male and female worms, especially in the organs involved in secretion and excretion. To test this, we determined the spatial localization of fucosylated and/or galactosylated proteins in the heads and tails of adult male and female *B. malayi*. We stained worms with both fluorescently tagged AAL lectin, which binds fucosylated LacNAc (enriched in males) and GS-I lectin, which binds to a-galactose residues (enriched in females). We observed a clear localization of fucosylated and galactosylated epitopes within the mouth and cephalic alae of both males and females consistent with the known distribution of sensory organs (**Fig. 5A**). Females showed a diffuse body staining with galactose-specific lectin GS-I, with higher levels at the cuticular areas and within the ovaries, compared to a much lower intensity of galactose staining across the body of male worms. In contrast, males displayed a very specific and high intensity fluorescence of fucosylated residues in reproductive organs and tissues (**Fig. 5B)** coupled to higher levels of fucosylated residues at the mouth and in the cephalic regions.

**Figure 5:**
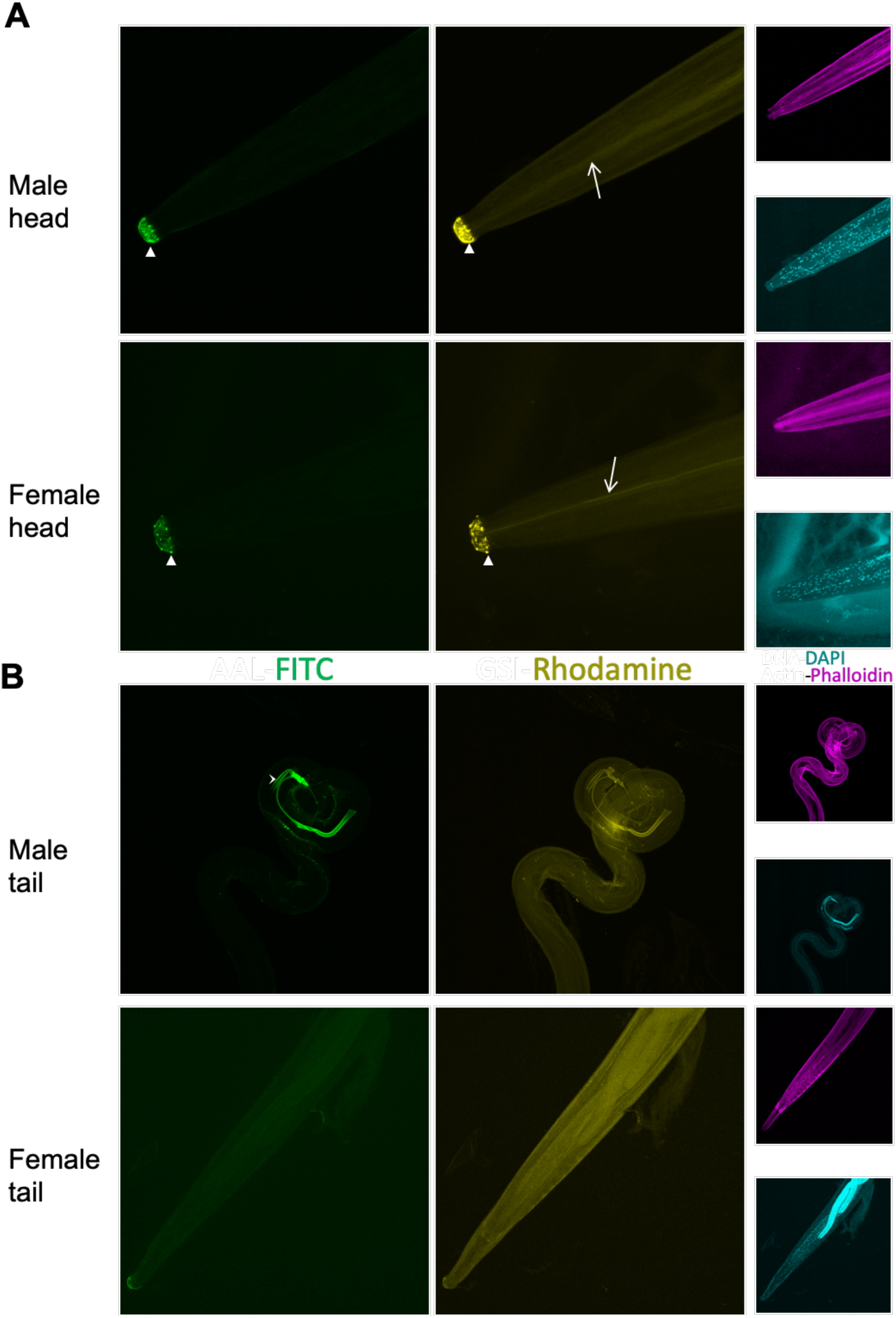
Fluorescence microscopy of male and female *B. malayi* adult worm heads and tails stained with fluorophore-labeled lectins. **A.** Heads of adult male and female worms. **B.** Tails of adult male and female worms. Images show full adult worms fixed in 4% formaldehyde 2:1 in heptane and stained with AAL-FITC for Fucosylated glycan epitopes (Green), GS1-Rhodamine for Galactosylated glycan epitopes (Yellow), Phalloidin for actin (Magenta) and DAPI for DNA (Cyan). Images are taken at 20x magnification for both males and females. Images shown are representative of multiple biological replicates (N=3). Arrow heads point to sensory organs in cephalic regions of both males and females, arrows highlight the intestines of both males and females and the pointy arrow indicates male spicules (reproductive organs).

### Adult male and female worms differentially glycosylate expressed proteins

The differential localization of galactosylated and fucosylated residues suggests such modifications occur on different proteins. To identify the protein partners of the sex-biased glycan epitopes, we carried out proteomic analysis on glycoproteins isolated with AAL and GS-I (**Fig. 6A**). Consistent with our findings from the lectin arrays, binding to AAL and GS-I showed higher affinity to male and female lysates, respectively (**Fig. 4**). Total protein extracts from both male and female worms were subjected to affinity chromatography on AAL and GS-I crosslinked columns. Chromatography fractions were checked for quality by silver stained SDS-PAGE gels (**Supp. Fig. 1**). Eluted proteins and column washes were labeled by tandem mass tags (TMT) and mass spectrometry analysis was done to identify the significantly enriched proteins in each fraction.

**Figure 6:**
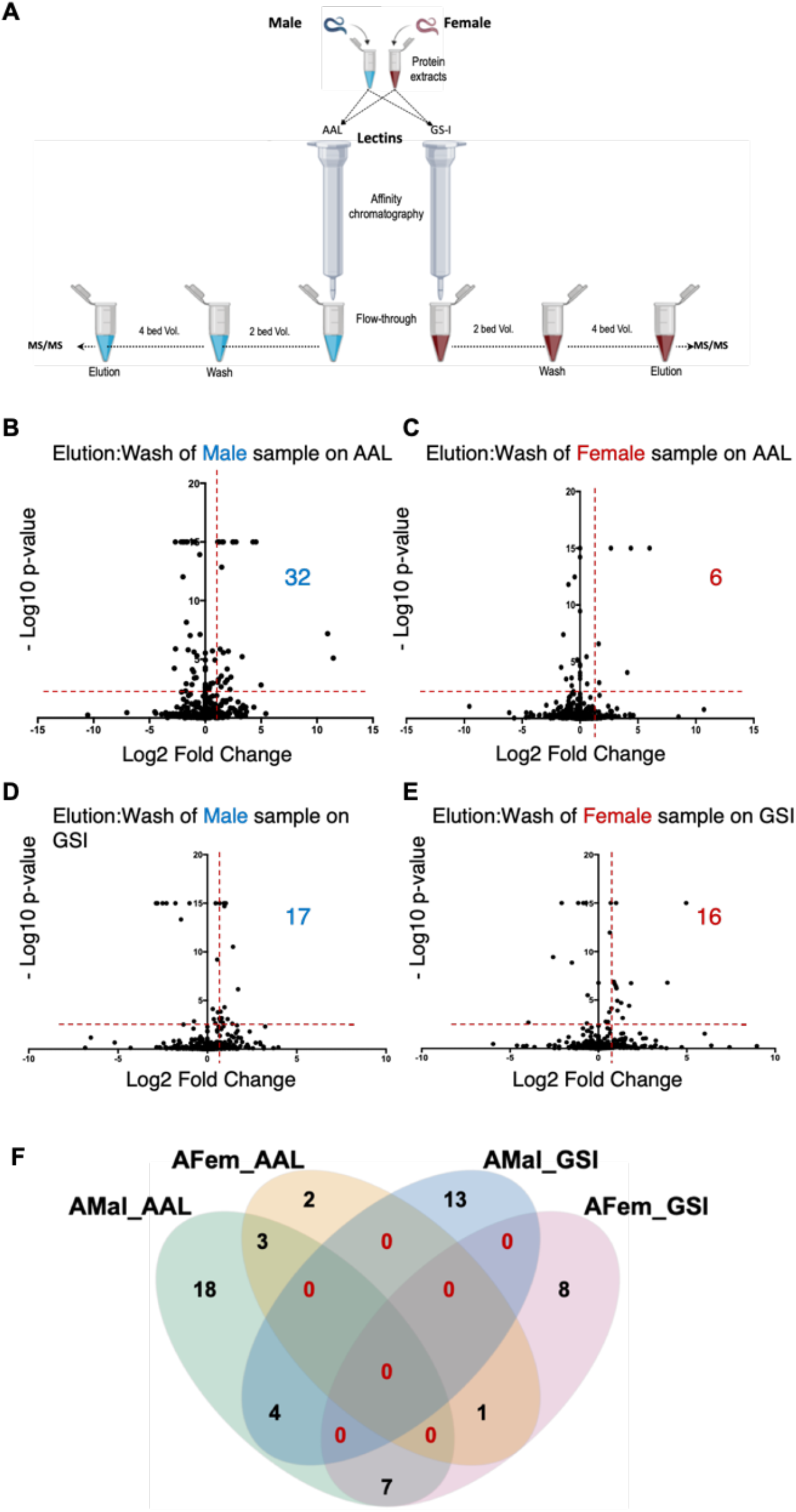
Mass spectrometry analysis of affinity chromatography fractions and glycoprotein characterization. **A.** Schematic diagram representing the experimental approach from total protein extraction to lectin affinity chromatography coupled with MS-MS. **B.** Volcano plot of MS/MS analysis data from full worm male lysate following affinity chromatography with GS1 lectin. **C.** Volcano plot of MS/MS analysis data from full worm female lysate following affinity chromatography with GS1 lectin. **D.** Volcano plot of MS/MS analysis data from full worm male lysate following affinity chromatography with AAL lectin. **E.** Volcano plot of MS/MS analysis data from full worm male lysate following affinity chromatography with AAL lectin. Plots show average Log2 of Fold Change (FC) of the detected protein intensity between wash and elution fractions across replicas with the corresponding - Log10 p-values. The total number of significantly eluted proteins from each fraction are shown with p-values <0.005 and at least 2 folds increase. **F.** Venn diagram of the overlap between the identified significantly eluted glycoproteins from both male and female samples on both lectins. Red numbers in the Venn diagram highlight the null intersections. malAAL: male worm lysate on AAL column. femAAL: Female lysate on AAL column, MalGS1: male lysate on GS1 column, FemGS1: female lysate on GS1 column.

Following statistical analysis and normalization (see Materials and Methods), we identified a total of 56 unique proteins significantly enriched in the eluate from either columns, out of ~ 950 proteins identified by MS/MS (two-way ANOVA, p < 0.005) (**Supp. Table S3a**). Of these proteins, 17 were galactosylated (bound to GS-I) and 32 were fucosylated (bound to AAL) in male worms, while 16 were galactosylated and only 6 were fucosylated in female worms (**Fig. 6B-D; Supp. Table S3b**). Clear sex-specific glycosylation of proteins was observed, with only 3/56 proteins displaying AAL-binding fucosylated epitopes in both sexes. **(Fig. 6F)**

To further investigate this subset of differentially glycosylated proteins, we profiled them for GO term enrichments (**Supp. Table S3c).** We observed a significant enrichment in proteins in the extracellular region (p-value = 0.0014). This finding, coupled to the detection of Bma-mif-1, a known immunomodulatory protein at the host-parasite interface, led us to determine whether these differentially glycosylated proteins were part of the *B. malayi* ES. All 56 detected proteins were observed in the *B. malayi* secretome defined in this study, suggesting a potential role for differential glycosylation in parasite glycoprotein-mediated immunomodulation.

### Guilt-by-association analysis reveals a cluster of highly secreted differentially glycosylated proteins with potential immune functions

Four secreted *B. malayi* proteins have previously been reported to be immunomodulatory: Bma-mif-1 (Prieto-Lafuente et al.), a macrophage migration inhibitory factor; Bma-far-1 (Zhan et al.), a fatty acid binding protein with stimulatory effects; Bma-CPI-2 (Manoury et al.), a cysteine protease inhibitor reported to be immunosuppressive; and the Cofactor Independent Phosphoglycerate Mutase Bma-IPGM-1 that induces a mixed Th1/Th2 response in the host (Singh et al.).

To identify immunomodulatory candidates in *B. malayi*’s secreted proteins, we made use of the guilt-by-association concept. We mined previously published *B. malayi* stage-specific transcriptomes for expression profiles of all ES protein coding genes and used dimensionality reduction (PCA, MDS) to identify genes that have similar expression profiles as the four known *B. malayi* immunomodulators (CPI2, MIF1, FAR1, and IPGM1). We clustered the results of both PCA and MDS analyses based on PC1:PC2 and X1:X2, respectively, to identify genes with the closest expression profile trends across all life stages. We then searched for known immunomodulators and identified the genes clustering with each of them. Both dimensionality reduction analyses and clustering (**Supp. Figure 2)** revealed a total of 112 proteins sharing similar expression profiles as the known immunomodulators, with more than 60% of those identified by both approaches **(Fig. 7A** and **Supp. Table S4)**.

**Figure 7:**
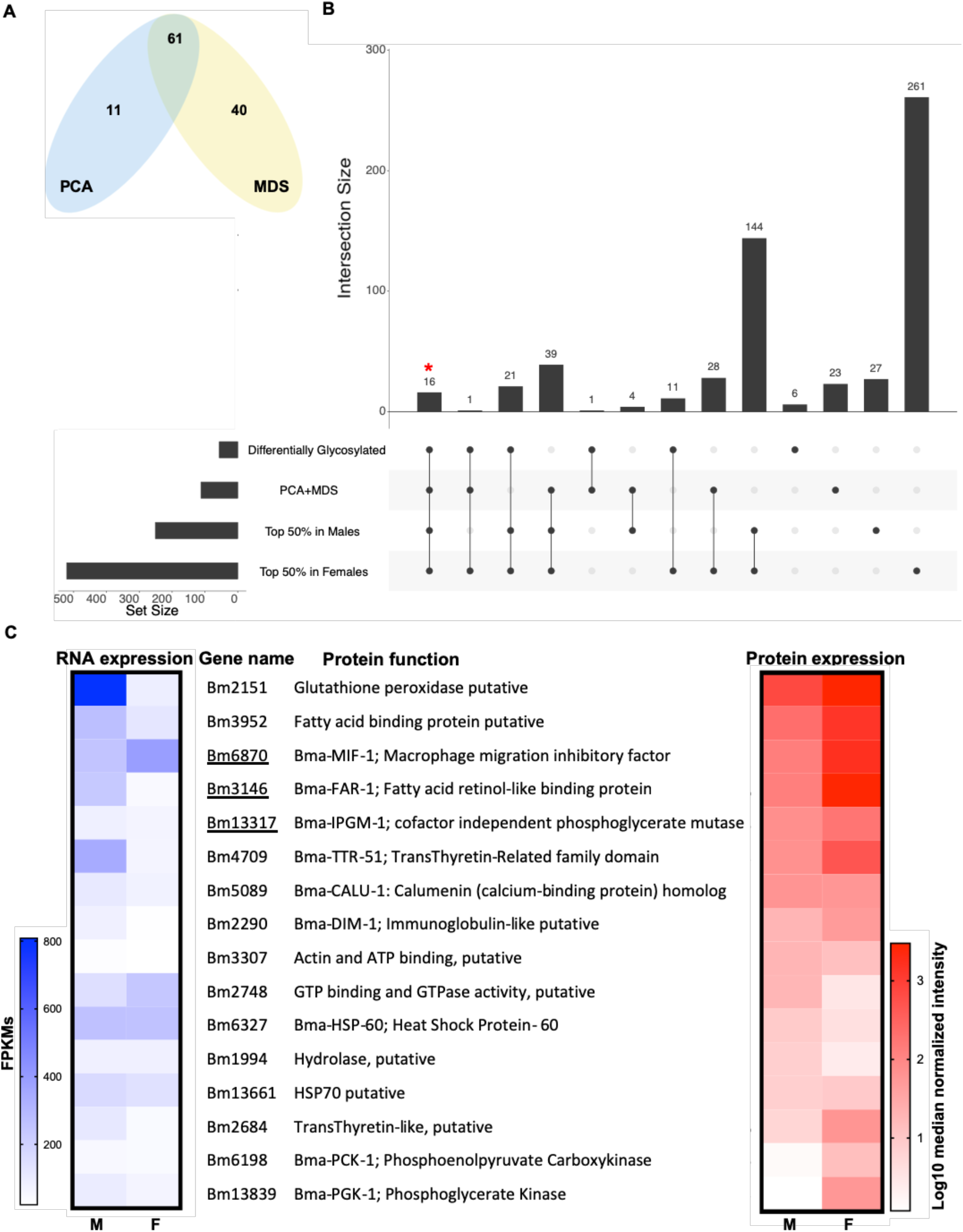
Immunomodulatory candidate proteins secreted and differentially glycosylated by adult male and female *B. malayi*. **A.** Venn diagram representing the overlap between immunomodulatory gene clusters following Principal Component Analysis (PCA) and Multi-Dimensional Scaling analysis (MDS) of stage and sex specific secretome transcriptomes. **B.** UpsetR plot representing the intersections across the candidate cluster genes, differentially glycosylated proteins and the top 50% proteins secreted from both Males and Females. **C**. Candidate proteins from intersections highlighted (**\***) in **B.** Gene names, protein functions and their corresponding RNA expression and protein intensity levels are shown. Highlighted by an underline are three previously characterized known immunomodulators.

To further shortlist the candidates that are highly expressed and differentially glycosylated by adult worms, we examined the overlap between the full set of differentially glycosylated proteins, the full set of *in silico* shortlisted candidates, and the top 50% secreted proteins detected in adult male and female worms. The analysis identified 16 proteins highly secreted by both males and females and differentially glycosylated, including three of the known immunomodulators secreted by *B. malayi*, Bma-MIF-1, Bma-FAR-2 and Bma-IPGM-1 **(Fig. 7B).** Gene identifiers, RNA expression levels, secreted protein levels, and individual function annotations for each of the 12 proteins are provided (**Fig. 7C**).

## Discussion

Protein glycosylation is an evolutionarily conserved posttranslational modification that plays an important role in multiple biological processes, from early development to sexual maturity of many living organisms (Varki, Ajit; Cummings, R. D.; Esko, J. D.; Freeze, H. H.; Stanley, P.; Bertozzi, C. R.; Hart, G. W.; Etzler and E.). The template-free synthesis of glycans leads to a large diversity in glycan epitope structures. Several such glycoproteins at the host-parasite interface in helminthic infections (McSorley et al.; Tundup et al.), including galectins, are known to regulate host-parasite interactions (Baum et al.; Boscher et al.). Glycoproteins also play crucial roles in sperm-egg biocommunication as well as immune signaling and activation (Giovampaola et al.). Characterization of the glycome is therefore an important component to gain a better understanding of the underlying biology of helminthic parasitism.

In this study, we profiled the glycoproteome of *B. malayi*, a model for filarial parasites. By analyzing male and female *B. malayi* transcriptomes separately for glycogenome expression, and experimentally probing their respective glycomes by lectin microarrays, we identified clear sex biases in glycogenome expression coupled to strikingly different lectin binding profiles between adult male and female *B. malayi*. The transcriptomic data suggested sex biases in glycan epitope expression, largely confirmed by lectin arrays.

Helminth glycomes are generally rich in oligomannose and paucimannose structures, extensively fucosylated, including the highly immunoactive α1-3 fucosylation, and rich in GlcNAc and GalNAc residues (Hokke and van Diepen). The lectin binding profiles show that *B. malayi* is no exception, and additionally uncovers a sex-biased fucosylation of type 2 LacNAc (LN) in males while females favor α1-2 fucosylation. Helminthic glycans may be branched with LacNAc and LacDiNAc residues, or otherwise truncated with simple terminal GlcNAc or GalNAc. Both *B. malayi* sexes expressed enzymes needed to catalyze the addition of β1,3-GalNAc in an O-linked manner to a Ser/Thr site forming the Tn antigen. The Tn antigen is a truncated glycan with terminal GalNac that is mostly known for its correlation with promoting metastatic colorectal cancers (Liu et al.). A higher affinity to core 1 and Tn-antigen binding lectins is reported in female worms as compared to male worms. This is partially explained by the higher expression of the ß-1,6-N-Acetylglucosaminyltransferases (Bm3610 and Bm5268) by males, catalyzing further modifications of the Tn epitope to core 2 and core 6 O-glycans. These epitopes are not detected by the core 1 O-glycan binding (AIA, MPA, MNA-G) and Tn-binding (HAA, HPA) lectins which show enhanced binding to glycoproteins from female worms.

To gain a better understanding of the sex bias in glycan epitope display, and identify any functional implications, we determined the distribution and localization of differentially expressed glycan epitopes in male and female worms. We observe the specific localization of galactosylated and fucosylated epitopes in reproductive tissues, sensory organs, intestines and cuticle. Such localization suggests a role for differential glycosylation in development, reproduction, and at the host-parasite interface, with sex specificity. To further investigate the functional implications of this differential glycosylation, we identified the protein partners of these differentially displayed epitopes. Using lectin affinity chromatography coupled to mass spectrometry we identified 56 differentially glycosylated proteins fucosylated and/or galactosylated in a sex-specific manner by *B. malayi*. The overrepresentation of secreted/excreted proteins amongst the differentially glycosylated proteins is not entirely unexpected yet the 100% overlap between ES proteins, as characterized in our study, and the differentially glycosylated proteins, strongly suggests a role for differential glycosylation in parasite interactions with its host. Substantial overlap with known secreted proteins (60%) validated the results.

The missing proteins identified in other studies (Bennuru et al) but not in our study may be due to sample variation and technological limitations. Most strikingly, we observed that most secreted proteins were shared across developmental stages, while previous work reported a stage-specific secretome (Bennuru et al.). This implies that the number of stage-specific secreted proteins may be much smaller than previously thought.

One of the most interesting differentially glycosylated, highly secreted *B. malayi* proteins is the well characterized Macrophage Migration Inhibitory Factor 1 (MIF1, Bm6870), a known immunomodulatory protein secreted by *B. malayi* (Prieto-Lafuente et al.) and determined in our study to be exclusively fucosylated in male worms. Fucosylation has been linked to CD14-dependent Toll Like Receptor 4 (TLR4) signaling activation (Iijima et al.). LeX, a fucose-containing glycan epitope, has also been implicated in mediating endocytosis of parasite glycoproteins to reach their effector sites. We hypothesize that fucosylation of MIF1 serves to decrease the immunogenicity of the parasite protein enhancing its suppressive function, or alternatively fucosylation may facilitate a higher internalization of the protein, similarly enhancing its suppressive function. An equally interesting differentially glycosylated protein is Bma-IPGM-1, previously shown to induce a mixed immune response, and which we showed displays a male-specific fucosylation coupled to female-specific galactosylation. These two well characterized *B. malayi* immunomodulators are joined by a third immunomodulatory protein Bma-FAR-1; two newly suggested drug targets, Bm5089 and Bm13839; and a subset of Heat Shock Proteins, Transthyretin-like proteins and a fatty-acid binding protein—all of which are part of protein families associated with host immunomodulation in helminthic infections, emphasizing possible sex-specific roles for *B. malayi* worms in the interactions with their hosts.

## Conclusion

In this study we show that adult male *B. malayi* preferentially display higher levels of fucose, localize the bulk of fucosylated and galactosylated glycoproteins to highly metabolically active internal and external reproductive organs, and differentially glycosylate secreted and immune-relevant proteins. Adult females favor α1-2 fucosylation (previously reported to play key roles in sperm-egg biocommunication), extensively galactosylate proteins, and localize the bulk of fucosylated and galactosylated glycoproteins to the intestines and secretory/excretory path with a clear signal from the anal pore and esophagus. Considering the respective localizations of fucosylated and galactosylated glycoproteins, and the differential glycosylation of secreted immunomodulatory and immune-relevant proteins, the data suggest that adult male and female *B. malayi* worms may be modulating the host immune response in sex-specific mechanisms and pathways. However, it remains to be determined if differential glycosylation of secreted proteins alters their effect on host cells. The immune potential of recombinantly expressed proteins with no glycosylation should first be established. The custom glycosylation of these recombinantly expressed proteins, if feasible, could help elucidate the functional implications of sex-dependent glycosylation of secreted proteins in *B. malayi*.

## Materials and Methods

### Worm culture and processing of the collected secretome

All *B. malayi* worms in this study were provided by the NIH/NIAID Filariasis Research Reagent Resource Center for distribution through BEI Resources, NIAID, NIH. A total of 30 adult males, 30 adult females, 300 L3, 300 L4 and 2×10^6^ microfilariae (mf) were used to characterize their respective secretomes. Adults were cultured in pairs at 1 worm/ml of RPMI1600 supplemented with 2% glucose. L3, L4 and mf were cultured in bulk in 5ml of RPMI1600 with 2% glucose. Spent media for all cultures was collected every 24hrs and filtered through 0.22μ filters, aliquoted and snap-frozen in an ethanol/dry ice bath. Collection of spent media was done for 7 days and combined aliquots were then concentrated 100x using Amicon Ultra15 with a 3Kda cutoff (cat# UFC900324) and buffer exchanged into cold sterile PBS, at pH 7.4. Concentrated samples were further split into 5 equal 100 μl fractions and methanol-chloroform precipitation of proteins was performed. Briefly, 400 μl of methanol was added to each 100 μl of protein sample and a subsequent 100 μl of chloroform was added on top. The mixture was then briefly vortexed and 300 μl of DI water added. Samples were then centrifuged at 13,000 rpm for 15 mins. Tubes were left at room temperature for another 15 mins to achieve a clear phase separation. The top layer was carefully removed leaving 50 μl of liquid. 400μl of methanol were then added and tubes tapped to induce mixing of phases. Samples were centrifuged as before, and the methanol wash step was repeated twice. Tubes were subjected to speed vacuum to evaporate all liquids and the resulting pellets were resolubilized using RapiGest SF Surfactant according to manufacturer’s protocol with a final concentration of 0.1% (RapiGest SF, 1 mg, 1/pk Part Number: 186001860). Samples were combined and analyzed by Mass Spectrometry.

### Glycogenome characterization and ortholog analysis

The full genome of *B. malayi* was downloaded from WormBase in 2018 and identified as B_malayi-4.0 assembly (Harris et al.).

#### Functional annotation

GO annotation of the genome was done using BLAST2GO v1.3.0 (Gö Tz et al.). A full list of protein glycosylation related GO terms was then obtained through the GO commission and can be found in **Supp. Table S5a**. The genome was filtered for genes associated with any of the glycosylation GO terms. A full list of 135 genes was identified and can be found in **Supp. Table S5b**. A similar analysis was done with the genome using the KEGG database (Kanehisa et al.) and all protein glycosylation related pathways can be found in **Supp. Table S5c**. Subsequently old gene identifiers from KEGG were assigned to the new IDs using BioMart for conversion (Smedley et al.). A total of 112 genes were found and are listed in **Supp. Table S5d**. To complement the functional analysis, *B. malayi*’s genome was analyzed using HMMER (Potter et al.) to annotate all Pfam domains. The full results are shown in **Supp. Table S5e**. Pfam annotations related to protein glycosylation were extracted from the Pfam database (Finn et al.); annotated genes can be found in **Supp. Table S5f**.

#### BLASTp analysis

A full list of glycosylation related proteins in human was extracted from Uniprot (D506-D515) and is shown in **Supp. Table S5g**. BLASTp was used to interrogate the whole *B. malayi* proteome against the compiled “glycosylation proteome”. Multiple blast hits were allowed and filtered for a maximum e-value of 0.05. All hits with at least 50% sequence identity were considered true and **Supp. Table S5h** includes all data from the Blastp analysis. Subsequently, hits with identity scores between 20-50% were further cross-checked against KEGG, GO and Pfam outcomes and hits present in at least one of the functional annotations were added. **Supp. Table S5i** shows a compiled list of unique genes per approach and the overall unique list.

#### Ortholog analysis

To evaluate the conservation of all glycogenes across several other nematode genomes, Biomart (Smedley et al.) was used and all available genome projects for the following organisms were selected and analyzed for orthologs and their percentage identity. Organisms selected were *Homo sapiens*, *C. elegans*, *F. hepatica*, *T. suis*, *O. volvulus*, *S. mansoni*, *D. immitis*, *W. bancrofti*, *B. pahangi*, *A. vitea* and *L. loa*. Full data are shown in **Supp. Table S5j.**

### Transcriptome sequencing and analysis

Lists of differentially expressed genes for male and female *B. malayi* were downloaded from (Grote et al.) and further filtered for glycogenome expression; **Supp. Table S2b** represents the full list of significantly differentially expressed glycosylation genes between adult male and female worms with corresponding functions. For dimensionality reduction, FPKM values used were generated as follows: BAM files were used with Cufflinks (v2.2.1) (Roberts et al.; Trapnell, Hendrickson, et al.; Trapnell, Williams, et al.) to obtain fragments per kilobase of exon per million fragments mapped (FPKMs) for each of the annotated transcripts and with Cuffnorm (Roberts et al.; Trapnell, Hendrickson, et al.; Trapnell, Williams, et al.) to obtain normalized FPKMs by library size. Data was filtered for secretome-coding genes as identified by our analysis. ClustVis (v1.0) (Metsalu and Vilo) was used for Principal Component Analysis and R was used to do Multi-Dimensional Scaling. PC1, PC2 and X1, X2 were then used for unsupervised clustering and plotting using the pHeatmap R package (https://cran.r-project.org/package=pheatmap, v 1.0.10).

### Protein extractions from male and female *B. malayi* adult worms

Protein extractions from both male and female worms were done by cryogenic grinding and PBS extraction. Briefly, worms were ground for 2 cycles of 10 mins in liquid N2 with 100 μl of PBS. The white powder was recovered and left to sit on ice until melted. Samples were then diluted 1:1 PBS and left overnight on an end-to-end rotator at 4°C. The samples were centrifuged for 15 mins at 13,200 rpm at 4°C. Supernatants were collected, quantified and aliquoted for further use. A total of 500 male worms and 200 female worms were used, split into two batches for biological replication.

### Lectin arrays

Lectin microarrays were generated as previously described (Pilobello, Krishnamoorthy, et al.). Briefly, arrays were manufactured in-house with a Nano-plotter v2.0 piezoelectric non-contact array printer (GeSiM) using a nano A-J tip. They were printed on Nexterion Slide H (Schott Nexterion) under 50% relative humidity at a surface temperature of 12°C. Commercial lectins and antibodies were purchased from Vector Labs, R&D Systems, Santa Cruz, TCI, AbCam, E.Y. Labs, or Sigma-Aldrich. The recombinant lectins rGRFT, rCVN, and rSVN were generous gifts from Dr. B. O’Keefe (NCI Frederick). A list of all printed lectins can be found in **Supp. Table S6a**. We note that we are unable to observe a subset of epitopes (e.g. α 2,8-linked sialic acids) in our array. Prior to sample hybridization, lectin microarray slides were blocked for 1 h with 50 mM ethanolamine in 50 mM sodium borate buffer (pH 8.8) and washed three times with 0.005% PBS-T (pH 7.4). Sample protein concentration and the degree of fluorescent label incorporation was determined by measuring absorbances at 280, 555, and 650 nm per the manufacturer’s instructions on a NanoDrop ND-2000c spectrophotometer (Thermo Scientific). A total of 5 μg of proteins per sample and contrasting labeled reference were mixed in 0.005% PBS-T (pH 7.4) for a final concentration of 100 ng/μL of protein. For reference, an equimolar mixture of all samples assayed was used. Slides were then loaded into a hybridization cassette (Arrayit) to isolate individual arrays (24 per slide). Samples were loaded onto individual arrays with a control array for the reference vs reference sample. Labeled protein samples were hybridized for 2 h at 25°C with gentle agitation. Following hybridization, samples were removed and arrays washed 4 times with 0.005% PBS-T (pH 7.4) for 10 minutes each. Slides were removed, submerged in ddH2O, and spun dry. Arrays were scanned using a GenePix 4300A array scanner (PMT 550 laser power 100% for both fluorescent channels). Background-subtracted median fluorescence intensities were extracted using GenePix Pro v7.2. Nonactive lectins were defined as having an average of both channel SNRs 90% of the data and removed prior to further analysis. Data were median-normalized in each fluorescent channel and the log2 of the sample/reference ratio was calculated for each technical replicate for each lectin. Technical replicates were then averaged for each lectin within each array. To identify significantly overrepresented glycan epitopes in male and female worms, a two-way ANOVA was done on the outcome of the lectin arrays and significant lectins and corresponding binding epitopes can be found in **Supp. Table S6b**.

### Lectin affinity chromatography

AAL and GSI lectins crosslinked to agarose beads were acquired from Vector lab (Cat# AL-1393-2), and EY lab (Cat# AK-2401-2) and packed into chromatography columns. Chromatography columns were used as per the manufacturer’s protocol. Briefly, columns were washed with 1.4M NaCl solution prior to use and re-equilibrated with PBS pH=7.4 before use. Samples were applied to the columns and flow-throughs collected by gravity flow. Columns were washed with 2 bed volumes of PBS and wash fractions were collected. Elution was done in 4 bed volumes with corresponding elution buffers (Glycoprotein Eluting Solution, Cat. No. ES-3100 Vector labs for AAL column and 0.1M Melibiose monohydrate for GSI Cat# AK-2401-2). Eluted samples were further concentrated using Amicon Ultra15 3Kda (Cat# UFC900324) to a final volume of 500μL (10x concentration). Elution buffer was exchanged to sterile cold PBS pH=7.4 and samples were aliquoted and stored for further use. Progression and quality of the chromatographies were assessed by SDS-PAGE silver-stained gels using Pierce^™^ Silver staining kit (ThermoFisher Scientific Cat# 24612) and standard 15% resolving gel 30:1 Polyacrylamide: Bis acrylamide denaturing gels. Gels were run for 1.5 hrs at 90V in 1x Tris-Glycine-SDS and Precision Plus Protein^™^ Kaleidoscope^™^ Prestained Protein Standards #1610375 was used for reference.

### Mass spectrometry analyses

To identify the proteins pulled down by the respective lectins, the last wash and the elution samples of each of the affinity chromatographies were subjected to mass spectrometry analysis. Similarly, we also analyzed resolubilized secretome samples. Samples were prepared as follows: Briefly, we denatured the proteins by heating for 15 min at 90°C. We added 1μg of mass spectrometry grade trypsin (Sigma Aldrich) and digested the proteins into peptides at 37°C overnight. For glycofractions, we measured resulting peptide concentrations with the Pierce Quantitative Fluorometric Peptide Assay (ThermoFisher, #90110); this step was not performed for the secretome samples. The glycofractions were labeled using TMT10plex Isobaric Label Reagent (ThermoFisher) while secretome samples were subjected to label-free analysis. The salt removal for both experiments was performed using Pierce™ C18 Spin Tips (Thermo Scientific, #84850) per manufacturer’s instructions. For both experiments, we used an EASY-nLC 1000 coupled on-line to a q-Exactive HF spectrometer (both Thermo Fisher Scientific). Buffer A (0.1% FA in water) and buffer B (80% acetonitrile, 0.5% acetic acid) were used as mobile phases for gradient separation. Separation was performed using a 50 cm x 75 μm i.d. PepMap C18 column (Thermo Fisher Scientific) packed with 2 μm, 100 Å particles and heated at 55°C. We used a 155 min segmented gradient of 0.1% FA (solvent A) and 80% ACN 0.1% FA (solvent B) at a flow rate of 250 nl/min as follows: 2 to 5 %B for 5 min, 5 to 25 %B for 110 min, 25-40 % B for 25 min, 49-80 % B for 5 min and 80-95% B for 5 min. Solvent B was held at 95% for another 5 min.

For label-free analysis of the secretome, the full MS scans were acquired with a resolution of 120,000, an AGC target of 3 x 10^6^, with a maximum ion time of 100 ms, and scan range of 375 to 1500 m/z. Following each full MS scan, data-dependent high-resolution HCD MS/MS spectra were acquired with a resolution of 30,000, AGC target of 2x 10^5^, maximum ion time of 150 ms, 1.5 m/z isolation window, fixed first mass of 100 m/z and NCE of 27 with centroid mode.

For the TMT labeled samples of the glycofractions, the full MS scans were acquired with a resolution of 120,000, an AGC target of 3e6, with a maximum ion time of 100 ms, and scan range of 375 to 1500 m/z. Following each full MS scan, data-dependent high-resolution HCD MS/MS spectra were acquired with a resolution of 60,000, AGC target of 2e5, maximum ion time of 100 ms, 1.2 m/z isolation window, fixed first mass of 100 m/z and NCE of 35 with centroid mode.

The RAW data files were processed using MaxQuant (version 1.6.1.0) to identify and quantify protein and peptide abundances. The spectra were matched against the *Brugia malayi* Uniprot database (downloaded August 18, 2018) with standard settings for peptide and protein identification, that allowed for 10 ppm tolerance, a posterior global false discovery rate (FDR) of 1% based on the reverse sequence of the mouse FASTA file, and up to two missed trypsin cleavages. We estimated protein abundance using iBAQ 3 for label-free experiments.

TMT labeled data was then filtered for proteins recovered only in both replicas for each sample type. Data is shown in **Supp. Table S3a.** Ratios of Elution:Wash and Wash:Elution were calculated for each detected protein by sample. Two-way ANOVA was performed using Prism v7 to identify significantly pulled down proteins in the eluted fractions as compared to their wash counterparts. An adjusted p-value cutoff of 0.05 and a minimum 2-fold change was used and a list of significantly eluted proteins can be found in **Supp. Table S3b**.

Label-free data relevant to stage and sex-specific secretomes were normalized by total intensities and median normalized by sample. Raw IBAQ values for all replicates and proteins are available in **Supp. Table S1a**.

### Lectin staining and confocal microscopy

Adult male and female worms were individually washed in PBS in a 12-well plate with 1ml of PBS per well. Worms were fixed and permeabilized in 4% paraformaldehyde 2:1 in heptane and left on high intensity shaking for 30 mins at room temperature. Fixed worms were washed three times in PBS for 5 mins and transferred to new 12-well plates with 1ml per well of 1x Phalloidin-iFluor 647 Reagent (Abcam cat# ab176759) in PBS pH=7.4 for 90 mins at room temperature with moderate shaking. Worms were washed in PBS three times for 5 mins and transferred to new 12 well plates with 1ml of Fluorescein labeled Aleuria Aurantia Lectin (Vector labs cat# FL-1391) and Rhodamine labeled Griffonia Simplicifolia Lectin I (Vector labs cat# RL-1102) in PBS pH7.4 with 0.2mM CaCl_2_ at 10 μg/ml final concentrations for both labeled lectins. Plates were left at 4°C overnight on moderate shaking and following that washed three times in PBS and transferred to new 12-well plates with fresh PBS. Unstained controls were processed similarly without the addition of the labeled lectins. Stained and control worms were mounted on microscopy slides in VECTASHIELD^®^ Hardset^™^ Antifade Mounting Medium with DAPI (Vector labs cat# H-1500) and left for 30 mins at room temperature before storing at 4°C until imaging.

All worms were imaged using a LSM Zeiss 880 confocal microscope with 20X air objective using the following settings: 1024 x 1024 pixels with 1.0 μm z stack step size, 8bit, 1.3 zoom. High magnification cut-outs were imaged using the 100X oil objective using 0.29 μm z stack step size and all other settings identical. Laser power was set at 1.2% (405nm), 2% (488nm), 2% (561nm), 4% (633nm). All tracks were imaged separately to minimize signal crosstalk. Projections were constructed using the ZEN2012 software with the “Maximum Intensity Projection” processing option. High magnification pictures were additionally filtered using the “Median Filter” processing option with x/y kernel size set at 3 voxels.

## Data Access

All raw glycomics data are made publicly available and can be found at doi:10.7303/syn24862046. The mass spectrometry proteomics data have been deposited to the ProteomeXchange Consortium via the PRIDE partner repository with the dataset identifier PXD024252.

## Acknowledgments

The authors would like to thank the NYUAD global PhD fellowship fund and the NYU Abu Dhabi Faculty Research Fund AD060 for supporting this work. This research was supported by funding from the Canada Excellence Research Chairs Program (LM). This work was also supported in part by the Division of Intramural Research (DIR) of the NIAID/NIH (EG).

## Supplementary Material

**Supp. Figure S1: Silver-stained SDS-PAGE analysis for chromatography quality control.** Molecular weight markers are shown, wash and elution fractions were loaded to evaluate the quality of the chromatography. A significant reduction in proteomic content is observed between the wash and elution fractions validating the chromatography outcomes.

**Supp. Figure S2: Dimensionality reduction and clustering of the *B. malayi* secretome**. A. MDS analysis of the *B. malayi* secretome coding gene expression across multiple life stages and sexes. Defined cluster represents the genes clustering with the known immunomodulatory protein coding genes (Highlighted in Blue). A single cluster of genes is reported to include all four known candidates. A total number of 101 genes are part of the cluster. B. PCA analysis of the *B. malayi* secretome coding gene expression across multiple life stages and sexes. Defined cluster represents the genes clustering with the known immunomodulatory protein coding genes (Highlighted in Blue). Two separate clusters are reported to include all four known candidates. Combined, both clusters include 72 genes clustering with known candidates.

**Supp. Table S1: Stage and sex-specific *B. malayi* secretome analysis.**

**Supp. Table S2: *B. malayi* glycotranscriptome analysis.**

**Supp. Table S3: Differential glycosylation comparative analysis between adult male and female *B. malayi*.**

**Supp. Table S4: Dimensionality reduction approaches to *B. malayi* ES transcriptome for immunomodulatory candidate identification.**

**Supp. Table S5: *B. malayi* glycogenome characterization by multiple functional annotation approaches.**

**Supp. Table S6: Lists of Lectins used for glycoprofiling of adult male and female *B. malayi* and corresponding list of lectins displaying sex-specific differential binding.**

