## Supplementary material for "Sex-specific glycosylation of secreted immunomodulatory proteins in the filarial nematode *Brugia malayi*": Supp_Figures

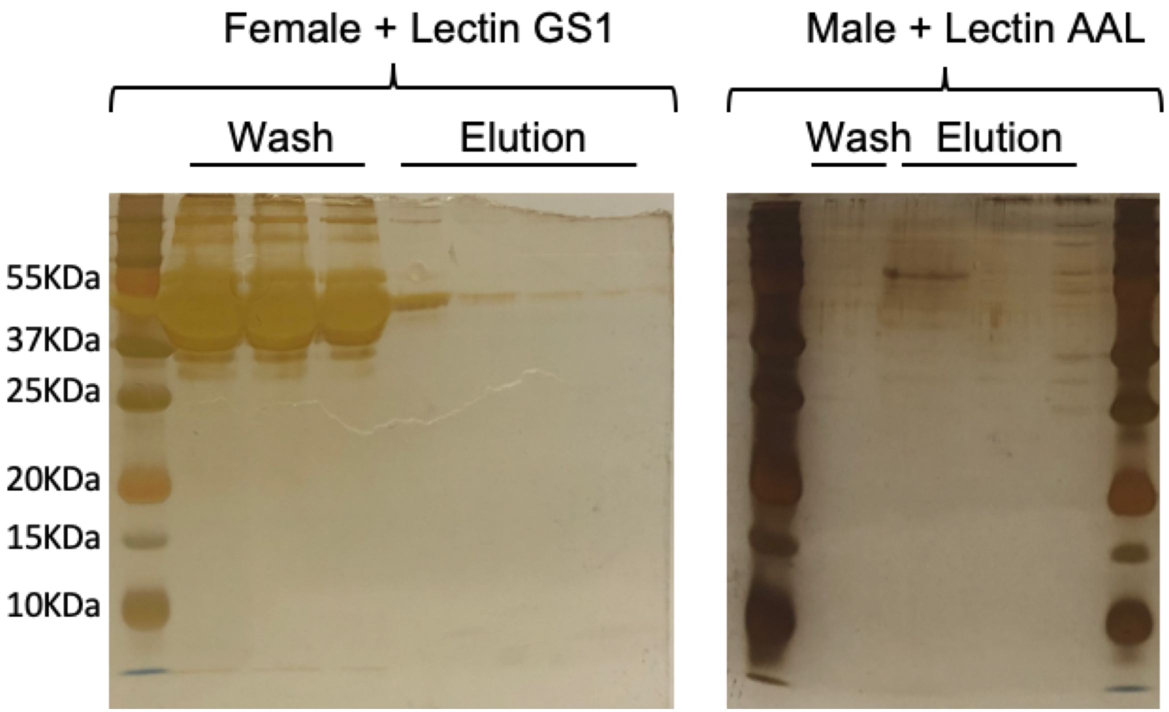


**Supp. Figure S1: Silver-stained SDS-PAGE analysis for chromatography quality control.** Molecular weight markers are shown, wash and elution fractions were loaded to evaluate the quality of the chromatography. A significant reduction in proteomic content is observed between the wash and elution fractions validating the chromatography outcomes.

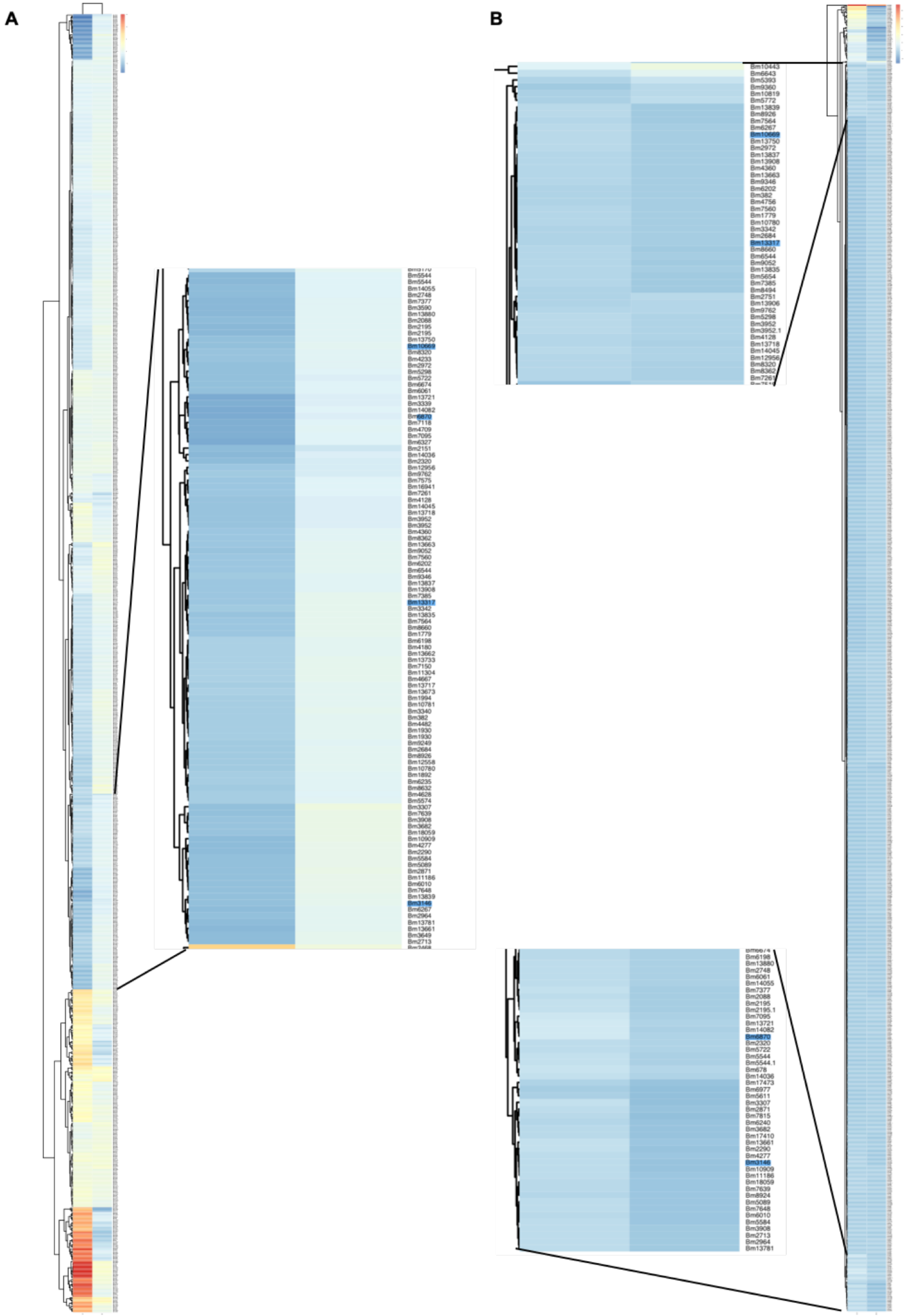


**Supp. Figure S2: Dimensionality reduction and clustering of the *B. malayi* secretome. A.** MDS analysis of the *B. malayi* secretome coding gene expression across multiple life stages and sexes. Defined cluster represents the genes clustering with the known immunomodulatory protein coding genes (Highlighted in Blue). A single cluster of genes is reported to include all four known candidates. A total number of 101 genes are part of the cluster. **B**. PCA analysis of the *B. malayi* secretome coding gene expression across multiple life stages and sexes. Defined cluster represents the genes clustering with the known immunomodulatory protein coding genes (Highlighted in Blue). Two separate clusters are reported to include all four known candidates. Combined, both clusters include 72 genes clustering with known candidates.
